# Optimizing data quality and completeness in visual proteomics experiments

**DOI:** 10.64898/2026.04.12.717927

**Authors:** Joseph M. Dobbs, Julia Mahamid

## Abstract

Cryo-electron tomography (cryo-ET) is fast developing from a tool primarily used to investigate structures of individual macromolecular complexes in situ into a high-resolution probe for molecular processes within diverse functional contexts in intact cells. It is thus increasingly necessary that the data are analyzed and quantified as completely as possible. But annotating and structurally characterizing macromolecular complexes with a high degree of completeness is a significant challenge, especially for smaller molecular targets. In particular, it is difficult to avoid incomplete localizations of complexes, false identifications, or losses during computational classification. To address these issues, we assessed parameters in data processing, including the role of voxel size in template matching, the effects of Volta phase plate imaging on localization, classification, and map refinement, and the extent to which multi-particle-based refinement of tiltseries improves these data processing steps. Our analyses provide practical guidelines that help maximize completeness in cellular cryo-ET data; accurate description of the sample is crucial for visual proteomics experiments, and these optimizations help ensure that data annotation and analysis are comprehensive.

## Introduction

While purified macromolecular complexes can be effectively localized in 2D single-particle cryo-electron microscopy (cryo-EM) images using a variety of approaches,^1–6^ it is generally far more difficult to localize targets in 3D volumes reconstructed from cryo-electron tomography (cryo-ET) data of crowded and heterogenous cellular samples.^7,8^ Losses incurred during particle picking or subsequent computational classification inevitably prevent accurate description of the sample. Such losses are expected in single-particle cryo-EM, where datasets of millions of particle images are commonly reduced to tens of thousands, either to isolate a rare state^9^ or to remove non-contributing particles damaged by factors including aggregation or denaturation at the air-water interface.^10,11^ In contrast, with cryo-ET data obtained from minimally perturbed cells, it is desirable to quantify as much of the cellular milieu as possible. Retaining the maximum number of complexes not only contributes to higher-resolution consensus reconstructions, but can contextualize the structure of interest through relationships between neighboring complexes,^12–15^ association with or proximity to other cellular structures or organelles,^16–19^ and correlations to cell state^20,21^ or cell type.^22,23^ However, this type of visual proteomics experiment^24,25^ represents a substantial challenge due to the complexity associated with analyzing such data at scale.

The ability to capture high-resolution structural information with cryo-EM was transformed by the introduction of direct electron detectors.^26^ An additional approach to increase the signal-to-noise ratio (SNR) and contrast of micrographs has been the use of phase plates,^27^ with the most practically useful of these to date being the carbon film-based, hole-free, Volta phase plate (VPP).^28^ However, VPPs induce unpredictable contrast-transfer-function (CTF) cut-on effects at low spatial frequency and produce an inconsistent phase shift.^29^ This can lead to difficulties in accurately estimating the CTF. Furthermore, VPPs appear to specifically attenuate high-resolution information through mechanisms that have yet to be fully characterized.^30^ Nonetheless, phase plates can be a valuable tool for boosting contrast in cellular cryo-ET data,^31^ especially when combined with recent developments in denoising,^32,33^ and signal^34^ or missing-wedge^35–38^ recovery. Still, the potential benefits of phase plates during the annotation and analysis of cellular data have largely not been systematically investigated.

Despite advances in deep learning-based particle localization,^6,39–42^ cross-correlation-based template matching^43–45^ remains the current gold standard for large-scale analysis of cryo-ET volumes.^12,15,46–49^ Traditional approaches have taken care to limit the introduction of high-resolution information which can lead to template bias in downstream structural analysis.^50^ However, recent work proposes that high-resolution information can improve cryo-ET template matching performance.^51^ Developments in 2D template matching in single, high-dose projections^52–54^ demonstrate that high-resolution information can indeed be effectively used,^55^ and given that projections and tilt series data are comparable in high-resolution information content,^56^ it stands to reason that a similar approach could also be useful in 3D. Nonetheless, it remains unclear to what extent highresolution information is retained in 3D tomograms.^57^

Tomogram reconstruction for high-resolution structural analysis is subject to numerous limitations, the most inhibitory being tilt-series misalignment^56^. Indeed, estimates of the resolution of cellular tomograms by Fourier Shell Correlation (FSC) are low, falling in the range of 20-50 Å.^58–60^ Thus, the contributions of high-resolution information in cryo-ET template matching require further clarification.

In cryo-ET data processing, much like in single-particle cryo-EM, computational classification follows particle localization. However, during the subtomogram analysis process, classification can fail to recover any, or all, of the particles of interest. Factors like the signal-to-noise ratio (SNR) of the images, the accuracy of tilt-series alignments, and the fraction of true positives in the data (driven by the quality of the initial localizations) impact the reliability of the classification procedure. In some cases, over 90% of annotated complexes of interest can be lost during classification.^49^ However, a clear understanding of which factors are limiting, and to what extent, is currently lacking. It is important to note that even a moderate 20% loss of complexes will have substantial impact on the quantitative analysis of cellular data (Fig. 1A): molecular concentrations would be underestimated, structural classes could be unequally lost, and particularly the spatial connectivity, *e*.*g*. of those ribosomes in polysome chains (Fig. 1B), may be substantially affected. These loses also affect, by reduction of particle numbers, the attainable final resolutions. Losses during particle picking and classification can therefore be severe impediments to describing the functional content of cells with cryo-ET.

**Figure 1:**
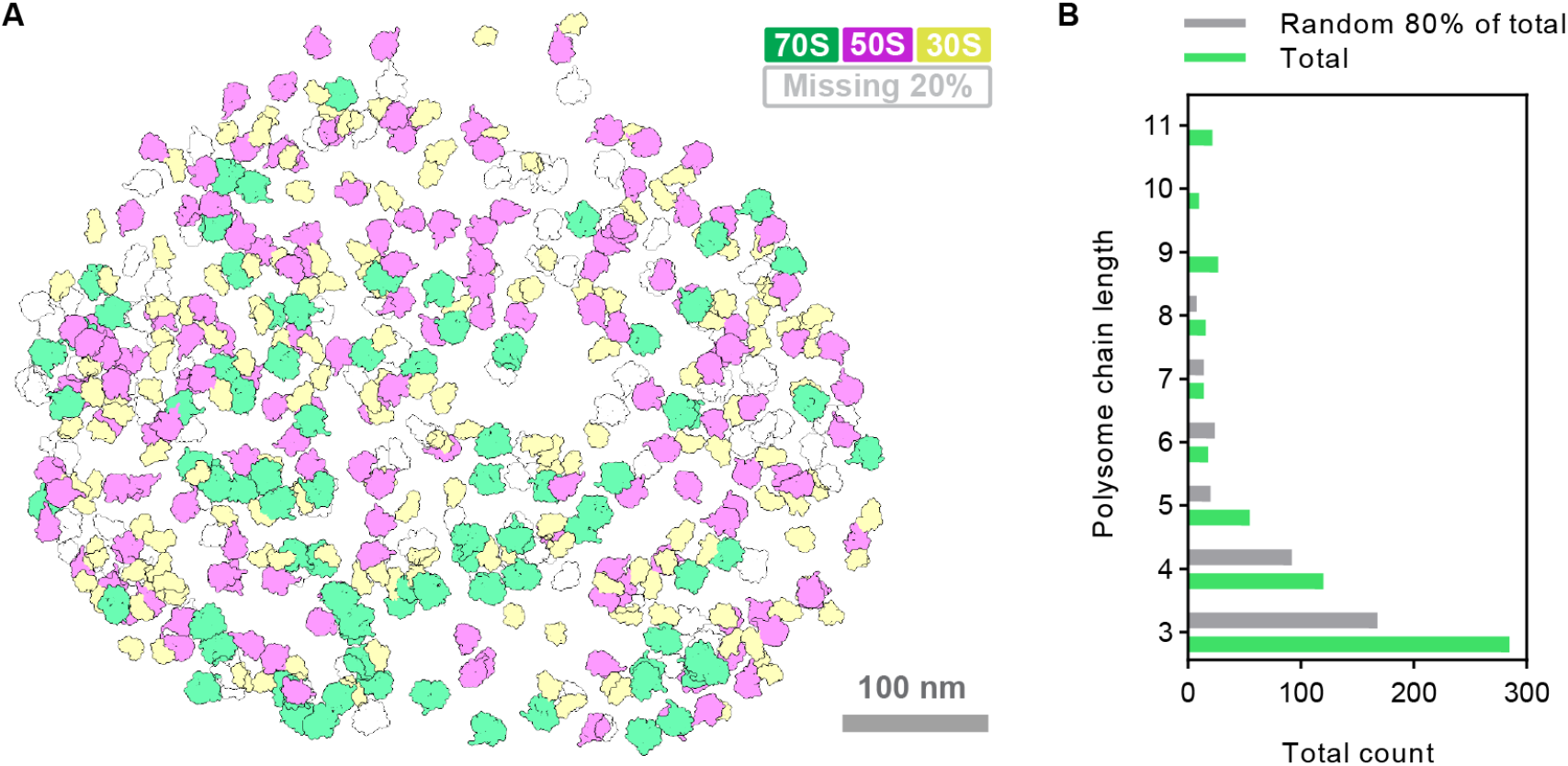
The impact of incomplete cellular cryo-ET data annotation on quantitative molecular description. **(A)** Representative mapping of 80% (by random selection) of manually curated 70S ribosomes, 50S large subunits, and 30S small subunits in one *Mycoplasma pneumoniae* cell. The missing 20% of annotations are transparent. **(B)** Representative quantitative analysis of the total counts of polysomes by chain length (number of 70S ribosomes) across 20 tomograms/cells, given complete annotation (green) and with 20% of ribosomes removed (grey). Polysome chain lengths were calculated by spatial distance analysis (7 nm cutoff between neighboring ribosomes, Methods). The small increase in 6-length polysomes is due to incorrect chain mapping introduced by the loss of a closer ribosome annotation.

Aiming to minimize such deleterious effects, here we systematically assessed the impact of multiple parameters, including use of the VPP, on the effectiveness of template matching, subtomogram classification, and map refinement. We show that searching cryo-ET volumes at smaller voxel sizes indeed improves template matching performance, but that high-resolution information has little to no effect. Our analysis highlights the importance of multi-particle based refinement of geometrical deformations and electron-optical parameters in tilt-series,^56^ not only for improving the final average map resolutions, but also during subtomogram classification, and especially for smaller targets. We quantified the VPP-induced loss of resolution for cellular complexes under typical cryo-ET acquisition conditions, and find that in accordance with expectations,^30,61^ map resolution indeed deteriorates by approximately 1 Å. Together, our results provide practical guidelines that help maximize completeness throughout cryo-ET data analysis.

## Results

### Establishing an evaluation dataset

Due to the difficulty in obtaining ground-truth data for quantitative evaluation of processing parameters, computational approaches in cryo-ET, especially for particle localization, have typically been assessed with lysates^62^ or simulated data,^63^ which largely do not recapitulate the complexity, crowdedness, and contrast of intact cellular samples. In this work, we took advantage of a subset of 20 high-quality and comprehensively annotated tomograms of *Mycoplasma pneumoniae* bacterial cells selected from our previous studies.^20^ These tilt-series (Table S1) capture single cells imaged at 1.7 Å/px with standard defocus (10 tomograms, Fig. 2A) or with a VPP plus defocus (10 tomograms, Fig. 2B). The tiltseries were aligned using 10 nm gold fiducials in IMOD,^64^ preprocessed and reconstructed in Warp (Methods).^65^

**Figure 2:**
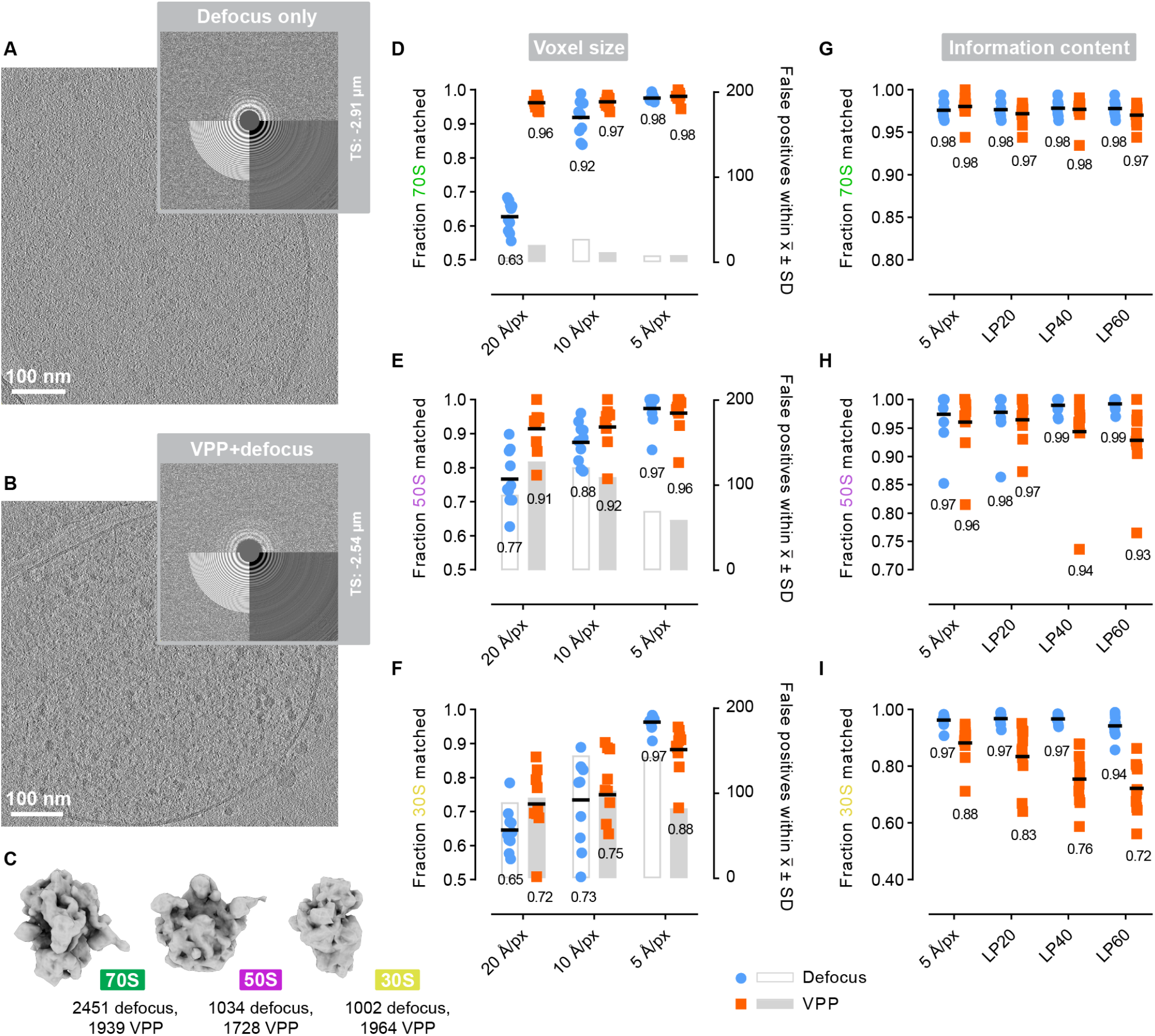
Template matching performance in cellular cryo-ET data improves with smaller voxel size. **(A, B)** Example slices through defocus-only (A) and VPP+defocus (B) tomograms of *M. pneumoniae* cells. Upper corner: power spectra computed with CTFFind5,^74^ and the estimated defocus for each of the tilt series (TS). **(C)** Subtomogram averages of the 70S, 50S and 30S derived from *M. pneumoniae* cells^20^ used for template matching (Methods). Averages are low-pass filtered to 10 Å for visualization. The total ground truth particle numbers in the 10 defocus-only and 10 VPP+defocus tomograms are provided. **(D-F)** Template matching performance at different voxel sizes. Dots represent tomograms, line and number represent mean. The left Y-axis (dots and mean value) indicates the fraction matched to annotated ground truth (true positives, Methods). The right Y-axis (bars) indicates the mean number of false positives with a score within one standard deviation (SD) from the mean 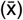 of the true positive cross-correlation values in that tomogram (at a cutoff of 400 peaks per tomogram). Analysis was performed for the 70S (D), the 50S (E), and the 30S (F). **(G-I)** The effect of low-pass filtering of the tomograms to different resolutions on template matching performance assessed by the fraction matched to annotated ground truth (true positives). Analysis was performed for the 70S (G), the 50S (H), and the 30S (I).

To interrogate the effects of various parameters, we chose three macromolecular targets of different sizes that are abundant and randomly distributed within the cellular volumes (Fig. 2C). The assembled 70S ribosome (2.5 MDa), which is commonly used to evaluate the effectiveness of particle picking,^40–42,51,66^ classification,^67–69^ and map refinement^56,70^ in cryo-ET data was chosen as a straightforward target: it can easily be localized with current particle picking tools due to its large size and dense RNA content. The free 50S large ribosomal subunit (1.6 MDa) can also be readily localized, albeit with more difficulty than the 70S, and was chosen as a medium-difficulty target. The free 30S small ribosomal subunit (0.9 MDa) was chosen as a more difficult target. Complete annotation of smaller complexes like the 30S in densely packed tomographic volumes is both a significant challenge and highly laborious, here requiring an exhaustive localization scheme combining traditional template matching, supervised deep learning, and extensive manual picking (Methods). Together, our comprehensive annotation of the 70S, 50S, and 30S complexes in intact *M. pneumoniae* cells (Table S1) establishes ground-truth data with realistic targets (at different levels of difficulty) for the testing and optimization of particle picking, classification, and map refinement.

### Evaluation of template matching parameters

Tomograms are typically reconstructed with voxel sizes larger than the original pixel size of the tilt images (“binning”).^43^ This has been necessary to keep file sizes within practical ranges, and to ensure particle localization algorithms like template matching (where running time scales inversely with voxel size) can be executed within a reasonable time on contemporary hardware. However, by necessity, binning entails removal of high-resolution information which has been reported to enhance template matching performance.^51^

We compared the performance of template matching for each of the three particle types in our defocus-only (Fig. 2A) and VPP+defocus tomograms (Fig. 2B) reconstructed at three different voxel sizes (20, 10, and 5 Å/px, Fig. 2D-F). While there are many implementations of template matching for cryo-ET,^43,45,51,65,66,71–73^ here we used the recently released pytom-match-pick^73^ (Methods). We assessed the raw template matching performance in terms of ground-truth recovery (within 100 Å tolerance, Methods) using the 400 highest cross-correlation scores per tomogram for each particle type. As a measure of the separation between true and false positives, we additionally quantified the number of false positives with cross-correlation scores that fall within one standard deviation from the mean for true positives per tomogram.

Overall, we found that template matching for each particle type in the VPP+defocus data recovered more true positives than in the defocus-only data at the larger voxel sizes, but performed slightly worse at 5 Å/px (Fig. 2D-F). Thus, the increased contrast from the VPP provided a boost to detection. However, a loss of performance becomes evident in the small voxel size tomograms, likely because per-tilt phase shift correction during template matching was not available in pytom-match-pick.^73^ Detection of 70S ribosomes was generally accurate, recovering >90% of true positives at 20, 10, and 5 Å/px, except surprisingly in the case of the defocus-only 20 Å/px tomograms (63% by mean). Detection of the 50S improved with smaller voxel size, reaching 97% recovery in 5 Å/px defocus-only tomograms. Similarly, 30S detection substantially improved at 5 Å/px, reaching 97% recovery in defocus-only tomograms, whereas only 73% were recovered at 10 Å/px. False positives were generally fewer in VPP+defocus data and at smaller voxel sizes (Fig. 2D-F). In searches for the 70S, false positives were uncommon (Fig. 2D), but were abundant for the 50S (Fig. 2E) and 30S (Fig. 2F), indicating a relatively poor ability to differentiate between true and false localizations for smaller targets. However, we note that false localizations to the abundant 70S complexes (which are made of 50S and 30S) were common in the 50S and 30S searches, particularly at smaller voxel sizes (Fig. S1). Together, these results indicate that template matching can indeed be improved by searching tomograms reconstructed at smaller voxel sizes, as previously suggested.^51^

However, the basis for this improvement remained unclear.^57^ We therefore examined how ablation of high-resolution features in the smaller-voxel tomograms affects template matching performance. We applied 20 Å, 40 Å, and 60 Å low-pass filters to the tomograms reconstructed at 5 Å/px using EMAN2^4^ and assessed ground-truth recovery when searching with templates of the 70S, 50S, and 30S (Fig. 2G-I). The templates themselves (5 Å/px) were not low-pass filtered to isolate the impact of high-resolution information in the reconstructions. We found that there was little, if any, loss of performance between unfiltered 5 Å/px tomograms and those low-pass filtered to 20 Å, 40 Å, and even to 60 Å (Fig. 2G-I), indicating that high-resolution features had no impact on the increased template matching performance at small voxel size (Fig. 2D-F). We next considered whether the actual representation of the structures of interest by a greater number of voxels drives this improvement. To test this, we upsampled the tomograms from 10 Å/px to 5 Å/px using pyTME’s density.resample function^66^ and assessed template matching performance (Fig. S2A). Ground truth recovery for any of the 70S, 50S, or 30S in the upsampled tomograms remained similar to that in the 10 Å/px tomograms, and did not increase to the level of recovery obtained for 5 Å/px tomograms (Fig. S2A). Furthermore, cross-correlation values of false positives (fig. S2B) were substantially higher in the upsampled tomograms (mean 0.093) compared to those reconstructed originally at 5 Å/px (mean 0.038). Thus, increasing the voxel counts did not contribute to the increased performance of template matching at smaller voxel size.

In applications of template matching on 2D projections, it has been demonstrated that interpolation errors from small differences in the assumed pixel size can have outsized impacts on the performance.^75^ Having empirically excluded the potential contributions of high-resolution information and greater voxel counts, we hypothesize that the improved performance is likely driven by the better representation of the entire spatial frequency range in smaller voxel reconstructions,^43,57^ leading to a reduction in positional errors which dampen localization peaks (see Discussion), and thus a better ability to distinguish true positives from the background.^73^ While here we do not consider theoretical aspects which might more completely describe the basis of our observations, in practice, our results support a particle localization strategy that leverages small voxel sizes, particularly for smaller molecular targets. However, the associated increase in computational cost underscores the utility of efficient template matching approaches^66^ for searching large volumes.

### Evaluation of tilt-series refinements in particle localization and classification

Resolution in 3D tomograms and subtomograms is critically affected by the accuracy of tilt-series alignments. In recent years, modeling of local sample deformations and refinement of misalignments and electron-optical parameters in tilt-series based on a constrained single-particle tomography approach^76^ has dramatically improved our ability to resolve structures to high resolution in cellular cryo-ET data.^56,70,77^ The program M expands on this concept and utilizes particle references to model the entire tomogram field of view.^56^ We reasoned that the benefits of geometrical and electron-optical refinement could then extend to improved template matching in the reconstructed tomograms,^78^ and enable greater performance during subtomogram classification (as previously suggested,^56^ but not empirically demonstrated).

We therefore applied M-refinement to all 20 tomograms in our evaluation dataset using the 70S ribosomes as reference particles. Overfitting to a single particle species was minimized by using conservative 2×2 image warping and 2×2×2×10 volume warping grids, in addition to CTF and pose refinements. Following M-refinement, structural details in the newly reconstructed tomograms appeared sharper (Fig. 3A, grey arrowheads). However, template matching performance at 5 Å/px for all three particle types was comparable to that performed on non-corrected tomograms (Fig. 3B, C). This result is in agreement with our finding that high-resolution features, despite improving during the tilt-series refinement process (as described below), are not effectively utilized in 3D template matching (Fig. 2G-I).

**Figure 3:**
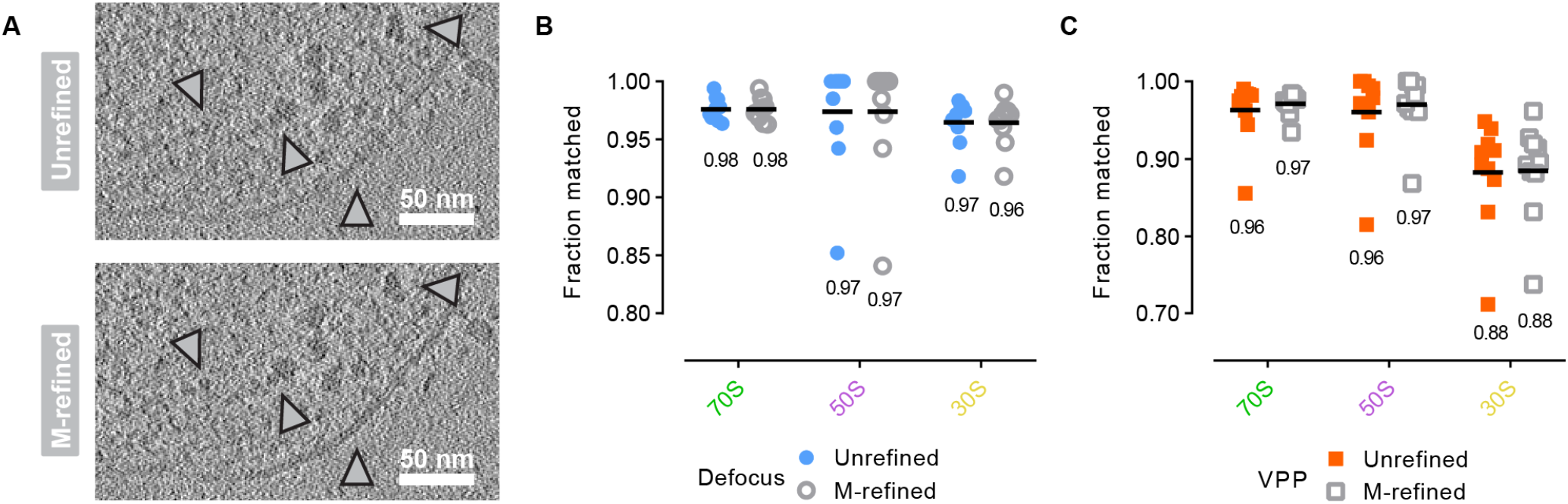
M-refinement improved details in tomograms, but did not affect template matching performance. **(A)** Tomographic slice through a VPP+defocus tomogram before (above) and after (below) M-refinement with 70S ribosomes (Methods). Structural details (indicated by grey arrowheads) appear sharper after M-refinement. **(B, C)** Template matching performance in unrefined and M-refined tomograms for the 70S, 50S, and 30S complexes. Analysis was performed in defocus-only (B) and VPP+defocus (C) tomograms reconstructed at 5 Å/px. Dots represent tomograms, line and number represent mean.

Next, we assessed the performance of computational classification of 3D subtomograms, before and after M-refinement, using RELION 4.0.^79^ We assessed defocus-only and VPP+defocus subtomograms reconstructed at 5 Å/px by including increasing proportions of decoy particles (non-target subtomograms, Methods), and quantified the fraction of the ground-truth particles recovered in manually selected “good” classes (Fig. 4A). Given the stochastic nature of particle classification,^1^ each experiment was performed in triplicate (Methods). Without M-refinement, classification performance decreased with a greater proportion of contaminating non-target subtomograms included in the dataset (Fig. 4B). Notably, for all particle types, VPP imaging-derived particles were more readily classified from non-targets than those from defocus-only data. We attribute this difference to the greater SNR in the VPP data.^80^ Crucially, we found that M-refinement improved the data quality such that classification was more effective across all particle types and imaging conditions (Fig. 4C). This was particularly evident for the smaller 30S: while in uncorrected defocus-only data 0% were recovered at 90% nontarget subtomograms, 94% were recovered when M-refinement was applied (Fig. 4B and 4C, right). Thus, reference-based refinement of tiltseries provides a substantial benefit to completely describing the contents of cellular cryo-ET volumes.

**Figure 4:**
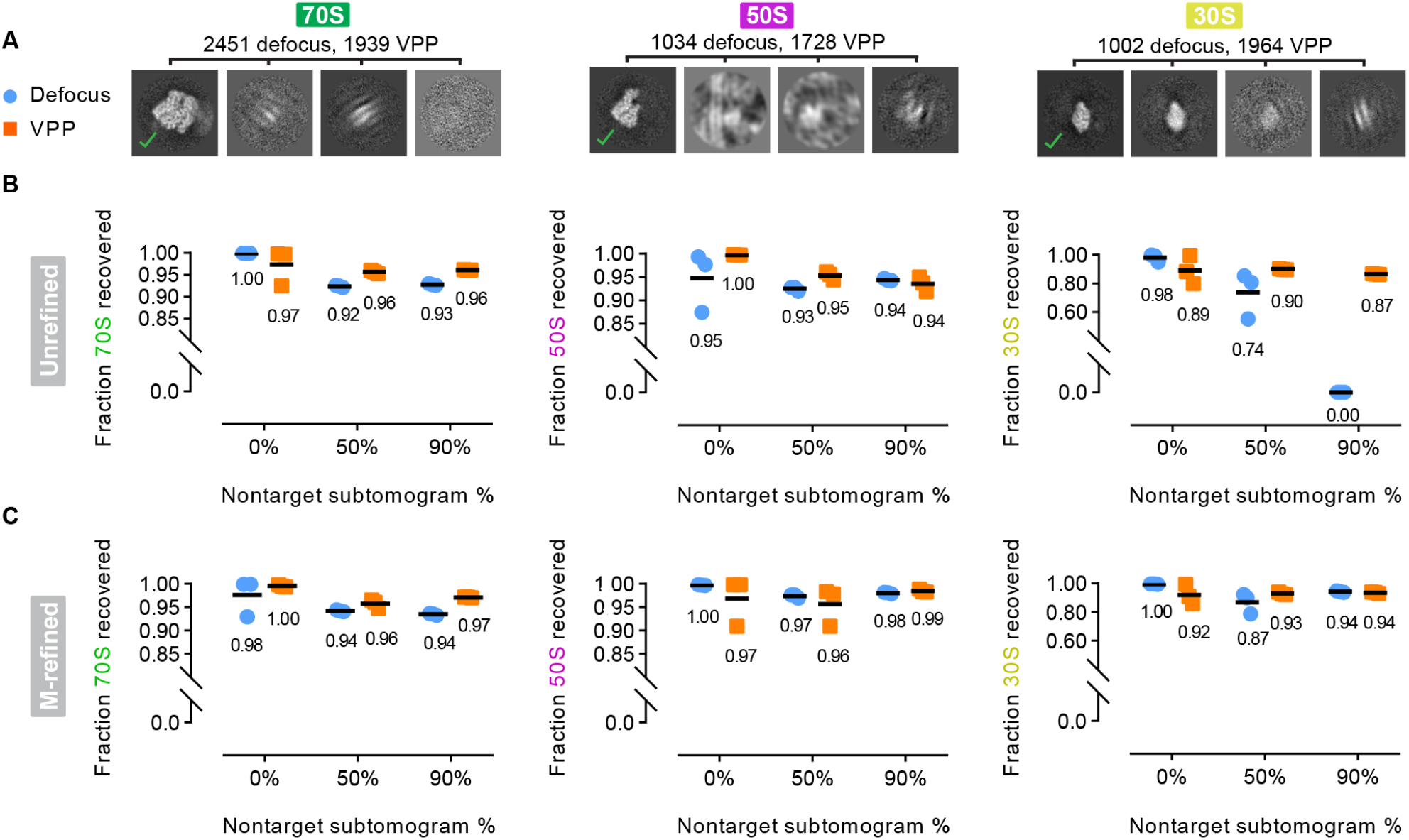
M-refinement with 70S ribosomes improves classification performance across particle species. **(A)** Example slices from RELION 3D classification: “good” classes are indicated by a checkmark. Analysis was performed with 70S (left), 50S (middle), and 30S (right) complexes. The total particle numbers in the 10 defocus-only (blue) and 10 VPP+defocus (orange) tomograms are indicated. **(B)** Fraction of ground-truth recovered through classification of subtomograms in uncorrected data, with increasing fractions of contaminating non-target subtomograms. **(C)** Fraction of ground truth recovered through classification of subtomograms in data refined in M with 70S ribosomes as references (Methods), with increasing fractions of non-target subtomograms. Dots represent classification runs, line and number represent mean.

### Evaluation of parameters influencing map resolution

Use of the VPP had appreciable benefits during particle localization (Fig. 2E-F) and in sorting target from non-target particles in subtomogram classification (Fig. 4B). However, use of the VPP is not recommended in single-particle cryo-EM experiments due to persistent loss of high spatial frequency information, leading to deteriorated final map resolutions compared to standard defocus-only imaging,^30^ and has been similarly counterindicated in cryo-ET based on analysis of symmetric virus-like particles.^61^ We aimed to quantify the scale of this loss under conditions common in cellular cryo-ET (*i*.*e*. lower particle numbers, thicker specimens, and asymmetric particles), where the increased contrast remains a useful benefit to visual interpretation.

With an expanded dataset of up to 10,000 subtomograms from defocus-only and VPP tomograms (dataset described in ^20^, Methods), we evaluated the attainable resolution for subsets of pre-aligned particles following a round of pose refinement in M. For the 70S ribosome, the global resolutions of maps generated from either 1,000 (10.3 Å defocus, 10.4 Å VPP) or 5,000 particles (6.7 Å defocus, 6.9 Å VPP) were similar in the two data types, although maps in the VPP+defocus data showed worse local resolution in multiple positions (Fig. 5A, grey arrowheads). This deterioration of high-resolution features was also apparent in voxel-based local resolution histograms (Fig. S3). In averages of 10,000 70S particles, the global resolution deteriorated from 5.5 Å in defocus-only data to 6 Å in VPP+defocus data (Fig. 5A). Indeed, Rosenthal-Henderson ResLog plots^81^ revealed a notably worse B-factor for VPP+defocus 70S averages (182 for defocus, 242 for VPP data; Fig. 5B). Similarly, the resolutions of 50S (Fig. 5C) and 30S (Fig. 5D) averages were approximately 1 Å and 0.5-1 Å worse in the VPP data, respectively. These results are consistent with previous benchmarking of viruslike particles^61^ and quantifications of VPP-induced deterioration of high spatial frequencies.^30^ They nevertheless demonstrate that obtaining maps with resolutions in the range of 6 Å from cellular cryo-ET data, even for smaller targets, is feasible with VPPs using a comparable number of particles to defocus-only data.

**Figure 5:**
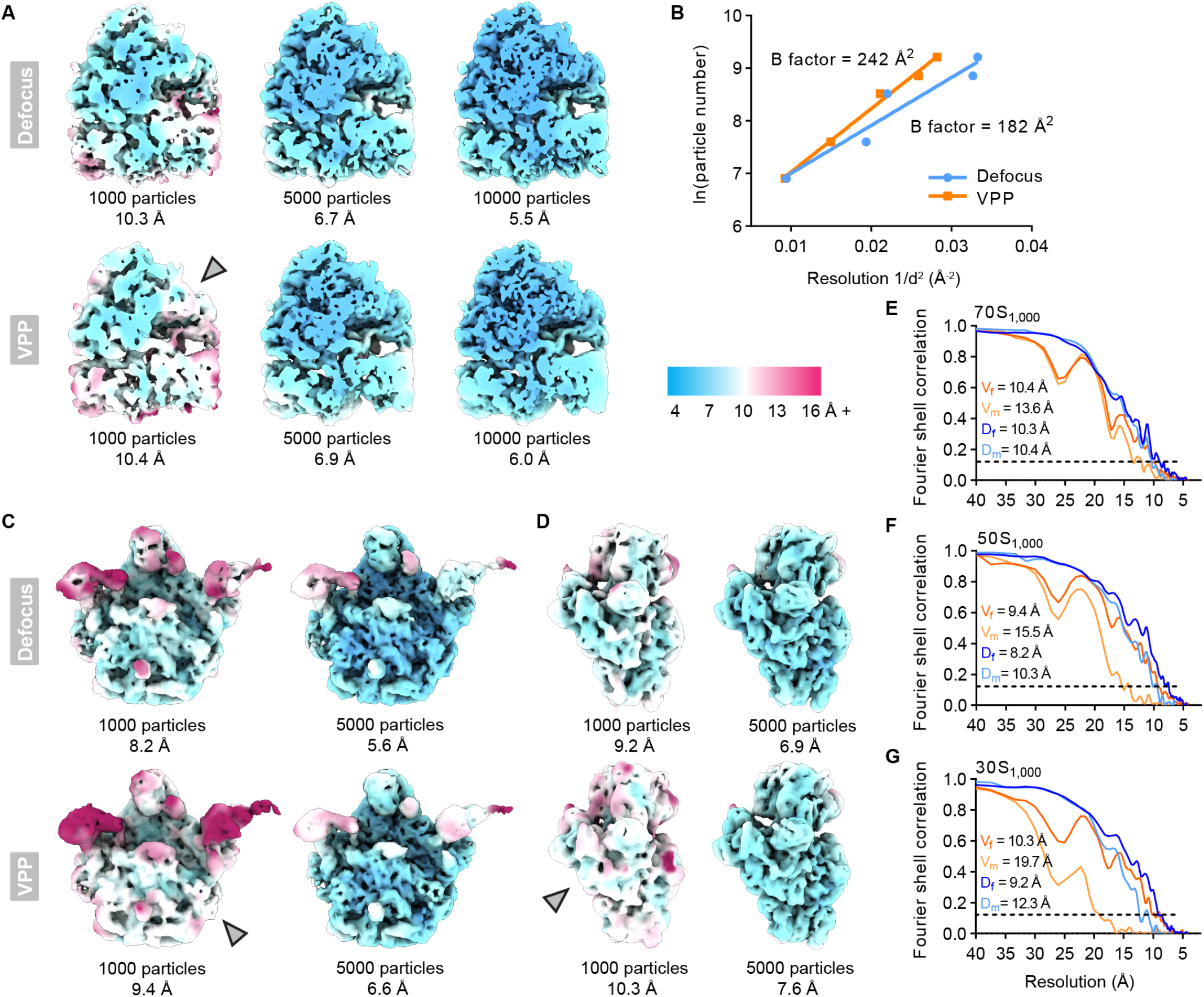
Effects of VPP use on attainable map resolution in cellular cryo-ET data. **(A)** Slices through defocus-only (top) and VPP+defocus (bottom) 70S maps generated from averaging different particle numbers (global resolutions based on FSC_0.143_ cutoff), coloured by local resolution (indicated by scale, right). Grey arrowheads indicate regions of lower map quality in VPP+defocus reconstructions as compared to defocus-only reconstructions. **(B)** Rosenthal-Henderson (ResLog) plot calculated for defocus-only (blue) and VPP+defocus (orange) 70S ribosomes. **(C, D)** Maps of defocus-only (top) and VPP+defocus (bottom) 50S (C) and 30S (D) generated from averaging different particle numbers, coloured by local resolution. **(E-G)** Fourier shell correlation curves for 1000-particle averages from defocus-only (D, blue) and VPP+defocus (V, orange) data under conditions of minimal (m) and full (f) M-refinement (Methods). Analysis was performed for 70S complexes (E), 50S complexes (F), and 30S complexes (G). In VPP+defocus data, the dip at around 27 Å was due to similar defocus values in the 10 analyzed tomograms (Table S1).

Lastly, we compared the effects of M-refinement with a minimal parameter set (single species only, image + volume warp) to M-refinement with a full parameter set (multi-species with the 70S, 50S, and 30S, image and volume warp, CTF refinement), to better evaluate the practical benefits of refining multiple parameters with a maximal number of reference particles (Fig. 5E-G). While the global resolution of averages from defocus-only data did not suffer from minimal refinement in the case of the 70S ribosome (Fig. 5E, 10.4 Å minimal vs. 10.3 Å full), full refinement was more important for smaller particle species (Fig. 5F, G): we found that the average of 1,000 50S particles reached a resolution 2 Å higher, and 1,000 30S particles reached 3 Å higher, when subjected to full refinement. In the case of VPP+defocus data, the effect was more pronounced: the global resolution of the 70S ribosome map was reduced by ∼3 Å with only minimal refinement (Fig. 5E), the 50S by ∼6 Å (Fig. 5F), and the 30S by ∼9.5 Å (Fig. 5G). Thus, utilizing the maximal realistic set of refinement parameters, and maximizing the number of reference complexes per tilt-series,^56^ are critical for corrections in both defocus-only and VPP+defocus data, especially when analyzing smaller targets.

## Discussion

Technology breakthroughs over the last decade now enable routine large-scale cryo-ET data generation^29,82–84^ from intact cells^85–88,49,89^ and multicellular samples.^90,91^ Realizing the full potential of this data requires that structures are analyzed with the highest possible degree of completeness, especially for experiments aiming to quantitatively describe and contextualize the functional states of a macromolecular complex, or a process, in cells. Here, we considered multiple factors that impact data analysis, and how these can be optimized to reduce losses during particle localization, through misclassification, and in final map refinements.

Our results support the notion that template matching performance is improved at smaller voxel size,^51^ and suggest that this may be of particular benefit for smaller complexes (Fig. 2F). However, our finding of nearly identical template matching performance in 5 Å/px tomograms when low-pass filtered to 20-60 Å (Fig. 2G-I) indicates that high-resolution structural features do not effectively contribute to localization in 3D data with current methods. Indeed, high-resolution information in cryo-ET data is constrained to the first few tilts of a series.^92^ Nevertheless, reconstructions at a smaller voxel size provide a more complete representation of the spatial frequency range described in the original tiltseries data. We suggest that the smaller voxels, by decreasing interpolations between pixels, reduce positional errors which dampen template-matching peaks and lead to better separation between true positives and the background.^73^ We hypothesize that the low-pass filters had little effect because the entire frequency range in the reconstructions was improved, and features ablated by the filters had no apparent impact. Our findings thus reconcile theoretical considerations of template matching in binned tomograms, where the shape and size of the template predominate,^57^ with practical assessments suggesting that performance can be increased at smaller voxel size.^51^ Appealingly, this suggests that the substantial benefits (Fig. 2D-F) of smaller-voxel template matching in 3D can be realized while avoiding the introduction of high-resolution template bias (by use of low-pass filtering). Computationally efficient^66^ approaches to template matching may therefore become especially useful for localizing complexes of interest.

While not currently a common choice for data acquisition, we aimed to systematically re-evaluate usage of the VPP during cryo-ET particle localization, classification, and map refinement. We show that VPP imaging provides a boost to performance when template matching is performed at voxel sizes of 10 Å/px or larger, while defocus-only imaging provides equal or better performance when tomograms are searched at 5 Å/px (Fig. 2D-F). The use of more accurate CTF information, including per-tilt phase shift correction, may in the future enable template matching in VPP data to match or surpass performance in defocus-only data. We also find that compared to defocus-only imaging, 3D classification in VPP+defocus data is more reliable, leading to higher recovery of targets in the dataset (Fig. 4B). The poorer performance in defocus-only data, however, can be improved (Fig. 4C) by refinement of tilt-series parameters^56^ with reference particles (here: 70S ribosomes). Following from the observed benefits of the VPP in template matching and in subtomogram classification, we quantified the scale of resolution loss associated with VPP imaging for maps of different complexes in our cellular cryo-ET data (Fig. 5A-D). In line with a previous study,^61^ we show that achieving resolutions in the range of 6-10 Å with the VPP is possible, but indeed comes with the penalty of an approximate 1 Å loss in global resolution. However, experiments on thin films indicate that the degradation of high spatial frequencies in VPP data^30^ makes reaching even higher resolutions increasingly difficult; by 5 Å, approximately 40% of the high-resolution signal present in the data has already been lost.^30^ In practice, the use of VPPs can be substantially complicated by several factors, including atypical optimal electron beam alignments^29^ and the variable quality of the VPPs themselves. Laser-based phase plates, in contrast, promise to provide reproducibly enhanced contrast with consistent phase shift,^93,94^ and without signal deterioration,^95^ but the effects of their use for cryo-ET remain to be investigated. Still, the increased contrast provides advantages for visual interpretation, particle picking,^40^ and when investigating rare or flexible assemblies where high-resolution reconstructions are infeasible^96^ but where segmentations may benefit.

Multi-reference-based refinement of tilt-series^76,70,56,67,77^ has dramatically improved the resolution of maps that can be obtained from cellular cryo-ET data. Here, we systematically assessed the potential benefits of tilt-series refinement beyond established practice, and particularly for particle localization and classification. Surprisingly, despite showing visual improvement to structural features in the 3D tomograms, we found that M-refinement had little impact on template matching performance in our evaluation dataset (Fig. 3B, C). We do not exclude the possibility that such refinements may improve template matching performance in other cases; it is possible that the robust gold fiducial-based tilt-series alignment procedure utilized here (Methods) prevented gross misalignments. Therefore, datasets where initial tilt-series alignments may be less robust, such as in FIB-lamellae data (currently), may benefit from multi-reference-based refinement to enable increased template matching performance. Crucially, we show that refinements based on abundant 70S ribosomes significantly improved the performance of 3D classifications, and extended to other (*i*.*e*. non-70S) complexes in the data (Fig. 4C). This was particularly effective for the recovery of smaller 50S and 30S complexes, and in the presence of a larger fraction of false-positive (non-target) subtomograms. Such improvements are especially useful given that obtaining complete annotations with 100% accuracy during particle picking is unlikely: this results in the common practice of particle over-picking and subsequent classification. Therefore, refining tilt-series using abundant macromolecular references, like ribosomes, to improve downstream classification performance will enable a more accurate and complete description of the sample. Finally, we show that multi-particle refinement utilizing the most complete set of parameters and references, via simultaneous refinement of multiple particle species^56^ and the CTF, leads to substantial improvement in the final map resolutions (Fig. 5E-G). This was particularly evident when a VPP was used in conjunction with defocus, and for smaller molecular targets.

In conclusion, this work informs complete and high-quality data analysis in cellular cryo-ET studies. Maximizing completeness has benefits both technically, where classification performance and map resolution are substantially improved by tilt-series refinement using abundant particle references, and biologically, where a visual proteomics approach^24^ can bridge structure and molecular sociology^8,97^ to contextualize macromolecular functions within cells.

## Acknowledgements

We are grateful to EMBL IT for HPC use, J. Bartho and S. Unger from the EMBL cryo-EM platform, E. Zagoriy and T. Hoffmann for technical support, R. Jensen and V. Maurer for insightful discussions, and L. Xue for data collection. We thank V. Maurer and D. Tegunov for critical feedback on the manuscript. J.M. acknowledges funding from the EMBL and a Chan Zuckerberg Initiative grant for visual proteomics (2021-234620).

## Author contributions

J.M.D. and J.M. conceptualized the study. J.M.D. designed experiments and performed analysis. J.M.D. and J.M. wrote the manuscript.

## Materials and Methods

### Cryo-ET data acquisition and tilt-series processing

Sample preparation, data acquisition and tilt-series processing were performed as detailed in previous work.^20,85^ Briefly, wild-type *M. pneumoniae* strain M129-B7 (ATCC 29342) were grown at 37° C in modified Hayflick medium.^98^ Quantifoil R2/1 200 mesh gold grids (Quantifoil Micro Tools) were plasma cleaned for 1 minute with a 90:10 argon:oxygen mixture (Fischione M1070) and UV irradiated for 30 minutes in a sterile laminar hood prior to cell seeding in 35 millimeter culture dishes. Vitrification was performed 16-20 hours after seeding using a manual plunger (manufactured by the Max Planck Institute of Biochemistry). Grids with adherent *M. pneumoniae* cells were briefly washed with phosphate buffered saline (PBS) containing 10 nm protein A-conjugated gold beads (Aurion), blotted from the back for 0.5-2 seconds, and plunged into a liquid ethane/propane mixture cooled by liquid-nitrogen.

Data acquisition was performed on a Titan Krios G3i (Thermo Fisher Scientific) transmission electron microscope operated at 300 kV, equipped with post-column energy filter (Gatan) and K2 direct electron detector (Gatan), using SerialEM software.^99^ Tilt-series were collected from -60° to +60° at 3° increment using the dose-symmetric tilt scheme^92^ and the following parameters: total fluence ∼120 e^-^/Å^2^, nominal magnification 88,000x, pixel size 1.71 Å, defocus range - 1.5 to -3.25 µm. For VPP data, phase shift was applied at approximately π/4 by conditioning each phase plate position for 90 seconds before the start of a tilt-series acquisition. A new phase plate position was used for each tomogram. For the larger dataset of subtomograms used in our map resolution analysis, additional tilt-series (reported in full in ^20^) were collected using the same microscope and settings, save for use of a K3 direct electron detector (Gatan), magnification (54000x) and pixel size (1.631 Å/px).

Tilt series were motion-corrected in SerialEM (defocus-only data),^99^ or Warp v1.09 (VPP+defocus data),^65^ CTF-corrected in Warp, aligned with IMOD eTomo^64^ via tracking of the 10-nm gold fiducials, and reimported into for Warp tomogram reconstruction. Tomograms were reconstructed at 5, 10, or 20 Å/px.

### Evaluation dataset

Particle annotations were constructed as extensively described in our previous work.^20^ Briefly, a total of 254 tomograms of *M. pneumoniae* were used in the analysis. Initial Particle references were obtained by manual picking in EMAN2^4^ (with 10 Å/px tomograms), followed by classification and refinement in RELION 4.0.1^79^ (with 5 Å/px subtomograms). The resulting averages were subsequently used for template matching in all 254 tomograms with PyTom v1.1.^45^ The resulting localizations were classified and averaged, and verified positions were used to train DeePiCt^40^ models for each of the three complex types. Localizations derived from these models were also classified and averaged, and combined with those from template matching. Finally, the combined particle lists were subjected to manual validation, removal of duplicates, and manual addition of missed putative complexes, in EMAN2.^4^ Multiple rounds of M-refinement^56^ based on all three particle species, followed by classification and averaging in RELION4,^79^ were performed. After three rounds of classification, there was no improvement in the number of complexes, and the complete list of particles was finally refined. The resulting map resolutions (FSC_0.143_ cutoff^100^) reached 3.8 Å for the 70S (51,496 particles), 4.2 Å for the 50S (27,655 particles), and 4.8 Å for the 30S (29,656 particles). These served as the references for evaluation of template matching performance in this work. The exhaustive localization scheme was necessary to maximize localization completeness for every target complex. A subset of 20 tomograms (Table S1: 10 with VPP+defocus, 10 with defocus only) was selected on the basis of their apparent quality, and complexes from these were used as the final ground-truth annotations for this study.

### Polysome analysis

Polysome analysis was performed with a custom python script (https://github.com/jmdobbs/cell_analysis_scripts/pyPolysome_count.py).^20^ The script measures the distance between a 70S ribosome’s mRNA exit site to the nearest 70S ribosome mRNA entry site. A 7 nm cutoff distance was selected, as in previous work,^20,46^ and the counts of ribosomes by polysome chain length were calculated.

### Evaluation of template matching parameters

Template matching was performed with pytom-match-pick v0.7.8^73^ using the data-derived consensus averages (described above) as references. Template matching was performed here at 7.5° angular sampling, with per-tilt defocus, dose weighing, a 500 Å high-pass filter, and random phase correction. A single phase shift value per tilt-series was used in the case of VPP+defocus data. Low-pass filtering of the tomograms was performed in EMAN2.^4^ The pyTME v0.3.3 density.resample function,^66^ with Fourier cropping enabled, was used to upsample 10 Å/px tomograms to 5 Å/px.

To assess the fraction of the ground-truth recovered, a custom python script (https://github.com/jmdobbs/cell_analysis_scripts/particle_identity_evaluator_v2.py) compared the positions of the top 400 cross-correlation score peaks per tomogram to the ground truth coordinates, with a 100 Å tolerance radius. This was performed for each of the three complexes. The script also measured the mean 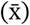 and standard deviation (SD) of the cross-correlation scores for the true positives (LCC_max_) and computed the number of false positives with scores within that range.

### Evaluation of classification parameters

Star files were constructed by adding non-target subtomograms to lists of true positive complexes. Coordinates for nontarget subtomograms were randomly generated from a list of manually selected points from inside the cells within positions where classification in RELION did not resolve any identifiable target. Subtomograms of all three target complexes and non-targets were reconstructed using Warp^65^ at 5 Å/px with a box size of 100^3^ pixels. Subtomograms were classified in 3D using RELION v4.0^79^ in single particle mode and the following parameters: spherical mask with diameter 400 Å, 10 classes, t=4, 7.5° angular sampling. Each classification used a reference for the 70S, 50S, or 30S derived from our previous work^20^, low-pass filtered to 50 Å. Each classification was performed in triplicate to ensure reproducibility. Following classification, “good” classes were manually selected and the fraction of ground truth particles recovered was assessed with the custom python script described above.

Classification parameter evaluation was performed with data from unrefined tomograms, where subtomograms were reconstructed using metadata created after tilt-series alignment in eTomo^64^, and with data from M-refined^56^ tomograms, where 70S ribosomes were used to improve alignments. M-refinement here was performed in 5 rounds with pose refinement, 2×2 image warp grid, 2×2×2×10 volume warp grid, and CTF refinement options.

### Evaluation of map refinement parameters

The impact of the VPP with regard to map resolution was assessed by comparing the resolutions and local map quality of defocus-only and VPP+defocus reconstructions derived from subsets of 70S, 50S, and 30S particles. Reconstructions were obtained by taking random subsets (1,000, 5,000, 10,000) using starparser^101^ of aligned particles from the consensus reconstructions and subjecting them to a round of pose refinement in M.^56^ For the 5,000 and 10,000 particle subsets, where the twenty tomograms that constitute our evaluation dataset in this work did not contain sufficient particles, additional subtomograms were obtained from additional VPP+defocus and defocus-only tomograms (254 in total) reported in our previous work^20^ and described above. The particles were reconstructed from fully M-refined tomograms (multi-species, pose refinement, 2×2 image warp, 2×2×2×10 volume warp, CTF refinement).

The impact of the parameter set used for M-refinement was assessed by comparing the resolutions of 1000-particle subsets of defocus and VPP data for each of the 70S, 50S, and 30S under conditions of full M-refinement (described above), or minimal M-refinement, where only geometric parameters (particle poses, 2×2 image warp, and 2×2×2×10 volume warp) were refined, and without multi-species correction (*i*.*e*. only for that particle type).

## Data and code availability

All data for the 20 tomograms of the evaluation dataset generated in this work, including raw frames or frame averages, tilt series, reconstructed tomograms and metadata, templates, and ground-truth STAR files, are deposited on EMPIAR (EMPIAR-13103). Subtomogram averages are deposited on the EMDB (defocus-only data: EMD-55551 to 55557, VPP+defocus data: EMD-55560 to 55566). The script used to assess ground-truth recovery in template matching is available on GitHub (https://github.com/jmdobbs/cell_analysis_scripts/particle_identity_evaluator_v2.py).

---

**Figure S1:**
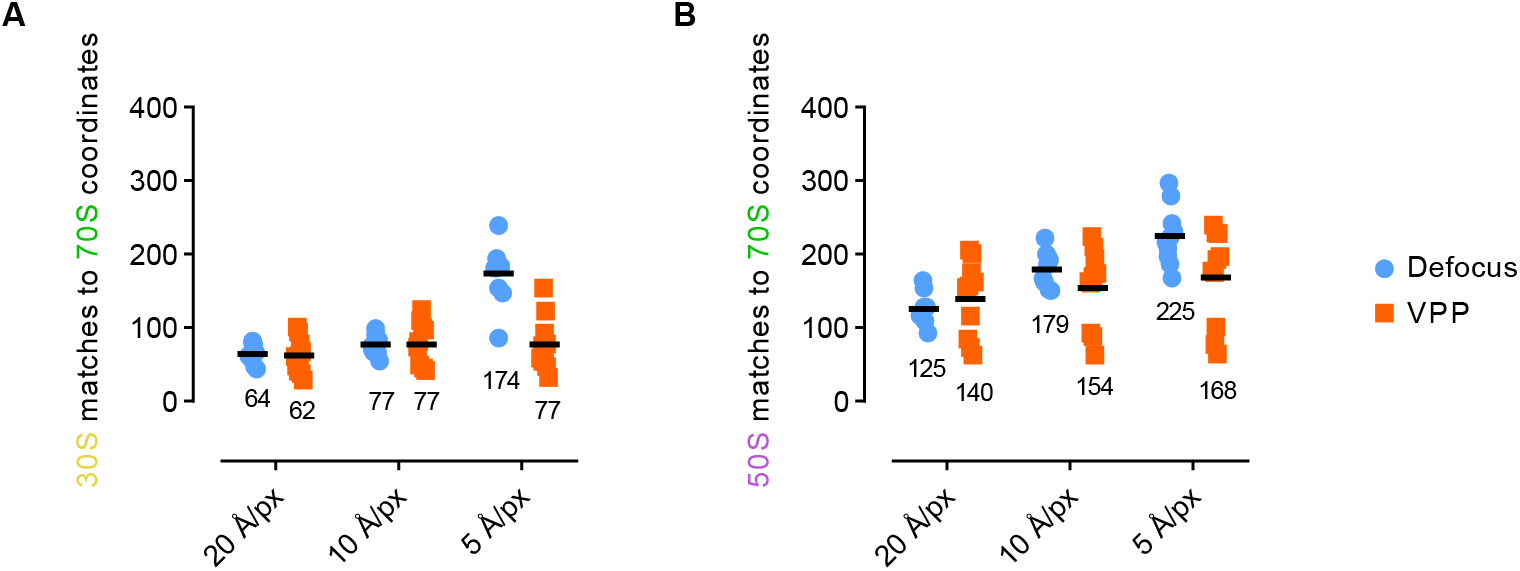
A substantial number of 30S and 50S localizations match to 70S coordinates, especially at small voxel size. **(A)** 30S localization peaks that match to true-positive 70S coordinates. **(B)** 50S localization peaks that match to true-positive 70S coordinates. Dots represent tomograms, line and number represent mean.

**Figure S2:**
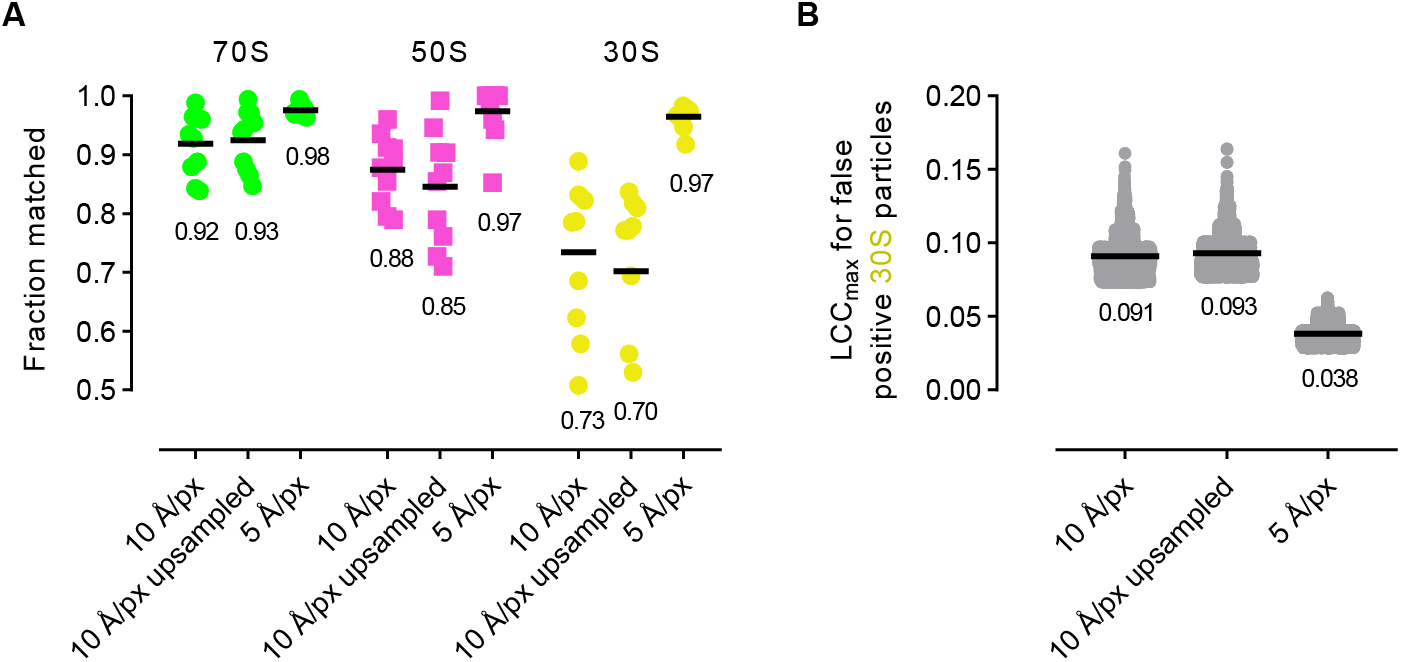
Upsampling 10 Å/px tomograms to 5 Å/px does not improve template matching performance. **(A)** Ground-truth recovery for template matching of 70S, 50S, and 30S complexes in defocus-only data for tomograms originally reconstructed at 10 Å/px, the same 10 Å/px tomograms upscaled to 5 Å/px (Methods), and the same tomograms originally reconstructed at 5 Å/px. Dots represent tomograms and lines and number represent mean. **(B)** Local cross-correlation maximum (LCC_max_) scoring results for false 30S identifications in defocus-only data, as described in (A).

**Figure S3:**
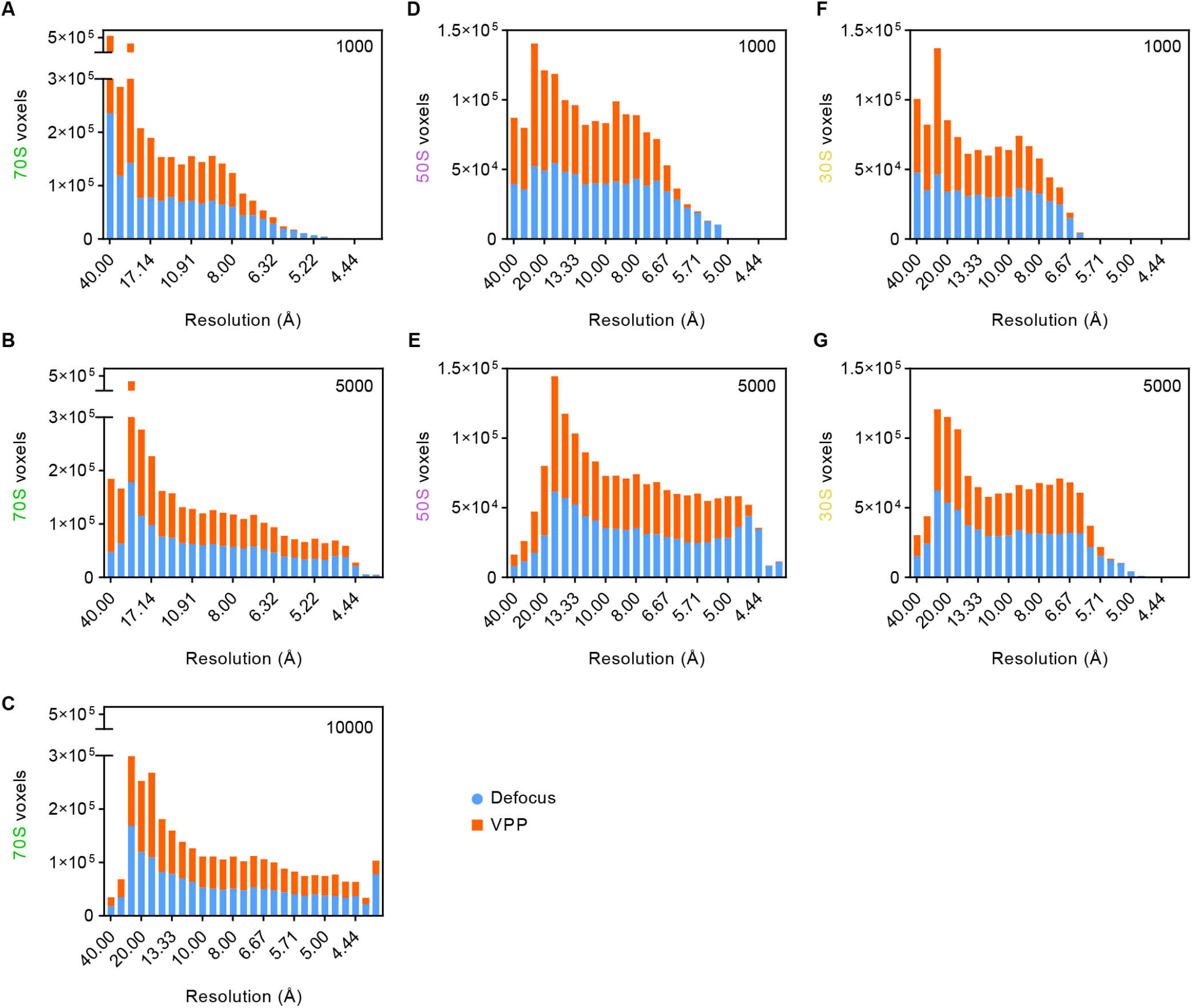
Stacked-bar local resolution histograms for 70S, 50S, and 30S complex reconstructions show a decrease in local resolution in VPP+defocus data. **(A-C)** Local resolution histograms of 70S ribosomes reconstructed from 1,000 particles (A), 5,000 particles (B), and 10,000 particles (C) from defocus-only (blue) and VPP+defocus data (orange). **(D-E)** Local resolution histograms of 50S reconstructions from 1,000 (D) and 5,000 particles (E) from defocus-only and VPP+defocus data. **(F-G)** Local resolution histograms of 30S reconstructions from 1,000 (F) and 5,000 particles (G) from defocus-only and VPP+defocus data.

**Table S1.** Detail of datasets and particle numbers analyzed in this work.

| Tomogram | VPP used | Phase shift ($\pi$) | Defocus ($\mu\text{m}$) | Total particles | Number of 70S complexes | Number of 50S complexes | Number of 30S complexes |
| --- | --- | --- | --- | --- | --- | --- | --- |
| c00035 | No | 0 | -2.03 | 546 | 263 | 156 | 127 |
| c00038 | No | 0 | -2.74 | 437 | 169 | 127 | 141 |
| c00041 | No | 0 | -3.40 | 582 | 256 | 177 | 149 |
| c00042 | No | 0 | -3.74 | 463 | 211 | 131 | 121 |
| c00080 | No | 0 | -2.91 | 342 | 194 | 69 | 79 |
| c00121 | No | 0 | -2.21 | 363 | 224 | 69 | 70 |
| c00208 | No | 0 | -2.15 | 422 | 304 | 57 | 61 |
| c00480 | No | 0 | -2.99 | 420 | 228 | 93 | 99 |
| c00498 | No | 0 | -1.56 | 539 | 353 | 88 | 98 |
| c00554 | No | 0 | -1.65 | 373 | 249 | 67 | 57 |
| aTS_30 | Yes | 0.25 | -2.65 | 504 | 107 | 184 | 213 |
| aTS_55 | Yes | 0.24 | -2.54 | 594 | 221 | 187 | 186 |
| aTS_56 | Yes | 0.24 | -2.63 | 640 | 215 | 202 | 223 |
| aTS_57 | Yes | 0.25 | -3.14 | 628 | 207 | 178 | 243 |
| aTS_60 | Yes | 0.18 | -2.51 | 533 | 240 | 158 | 135 |
| aTS_61 | Yes | 0.24 | -2.92 | 584 | 242 | 153 | 189 |
| aTS_65 | Yes | 0.21 | -3.29 | 920 | 160 | 379 | 381 |
| aTS_66 | Yes | 0.21 | -3.35 | 243 | 65 | 63 | 115 |
| aTS_68 | Yes | 0.17 | -2.54 | 480 | 244 | 112 | 124 |
| aTS_69 | Yes | 0.19 | -2.94 | 505 | 238 | 112 | 155 |

